# Convergent and divergent neural and behavioral responses to chronic cocaine use in macaques

**DOI:** 10.1101/2025.09.23.678075

**Authors:** Ana M.G. Manea, Tait Erickson, Gabriela Delgado Salazar, Sarah R. Heilbronner, Anna Zilverstand, Jan Zimmermann

## Abstract

Chronic cocaine use leads to physical dependence, psychological addiction, and various health problems such as cardiovascular issues and cognitive impairments. This condition continues to be extremely difficult to treat, as we grapple with its neural and behavioral complexities. What we are missing is a comprehensive account on how naïve brains react to occasional and chronic cocaine use, abstinence and re-exposure within subject. We tracked naïve macaques using precision functional mapping before, during and after chronic cocaine self-administration over a period of nine months. We analyzed the entire brain to illustrate the pervasive impact of chronic cocaine use on overall function. Although we set out to map the “addiction” connectome, we found that the neural correlates of chronic drug use and recovery are more complex that can be captured by a singular, canonical connectome. We provide a preclinical model that maps the progression of the macaque functional connectome from a naïve state through chronic cocaine use, recovery and re-exposure. We demonstrate that the “addiction” connectome is dynamic and is characterized by the interaction of subject by drug by time. Occasional cocaine use, whether initial or re-exposure, overrides individual differences, leading to a strong homogenous effect across subjects, or convergence. However, the brain’s response to prolonged cocaine use becomes more individualized, resulting in the divergence of effects across subjects. This can be explained by heterogeneity in cocaine-induced effects and in their associated timescales. Abstinence-induced neural alterations converged across subjects after 1-month, but by 2 months, individual trajectories revealed heterogeneity, possibly indicating differences in recovery.

## Introduction

Substance use disorders (SUDs) remain a pervasive global challenge with significant socio-economic and health implications, affecting millions worldwide^12^. While treatments are available, long-term relapse rates, even for those who do start treatment, are staggeringly high at up to 85%^3–5^. This reality underscores the urgent need for a better understanding of how SUDs are initiated and progress. Primarily, this dearth of knowledge arises from the challenges of implementing longitudinal studies and the ethical and feasibility limitations on experimental studies in humans. In contrast, animal models enable tracking within-subject neurobehavioral changes from baseline through chronic drug use, abstinence and re-exposure. Hence, prioritizing and continuously developing better and more ecologically relevant preclinical animal models is crucial for elucidating the neural and behavioral changes associated with exposure to and abuse of addictive drugs. Moreover, resting-state functional magnetic resonance imaging (rs-fMRI) studies in individuals with SUDs, regardless of substance, have revealed altered connectivity across all major large-scale brain networks ^6–10^. Thus, it is critical that a preclinical animal model of SUDs enable the acquisition of whole-brain data, with rs-fMRI being particularly translatable across species.

Nonhuman primates (NHPs) offer distinct and appealing characteristics to study the neurobiology of drug seeking and taking^11,12^. First, their phylogenetic proximity to humans results in similar brain organization and function^13–15^. Second, NHPs demonstrate similar drug biodistribution, pharmacokinetic and pharmacodynamic profiles to humans^19,2019,2016–18^. Finally, NHP models of chronic drug self-administration (SA) demonstrate strong predictive and construct validity regarding compulsive drug use in humans ^19^. Overall, NHPs have significant translational value^11,20^. Hence, combining NHP models with ultrahigh-field rs-fMRI^21–23^ promises to significantly improve the ecological validity and translatability of findings from animal models.

To date, progress in translating rs-fMRI methods into clinical practice have been limited ^24^. Part of the reason has been our limited data quality (signal-to-noise), quantity and fidelity (contrast-to-noise) within individual subjects, requiring averaging over many subjects to achieve statistical significance. In turn, this leads to a lack of single-subject reliability which hinders our efforts to achieve individual-specific characterization of brain-based disease processes. Precision neuroimaging, however, has emerged as a transformative tool, allowing us to uncover functional dynamics in unprecedented detail. First developed for human neuroimaging, applying precision neuroimaging in animal models promises to enhance our understanding of the timescales of SUD progression. Coupled with within-subject longitudinal designs, it also addresses the problem of inadequate matching between control and case samples in group comparisons. A case-control group-level analysis treats interindividual differences in brain organization and how the brain reacts to addictive drugs as noise, which can hinder efforts to develop a brain-based model of SUDs by producing blurred or misleading results. It is widely recognized that neurobiological interindividual differences are reliable and are behaviorally relevant^24–29^. Precision neuroimaging accounts for interindividual differences by collecting dense high-quality measurements through repeated sampling of the same individual. Here, we set out to demonstrate that interindividual differences extend beyond variations in brain organization and how subjects respond to chronic cocaine SA, encompassing differences in the timescales of disease progression as well.

To achieve these goals, we acquired very long sessions (70 mins) of high-quality ultrahigh-field rs-fMRI (10.5 Tesla) at nine time points in four naive macaque monkeys from baseline to chronic cocaine use, abstinence and re-exposure. To further increase translation, we employed an SA paradigm, the gold standard in preclinical substance dependence research, due to its similarity to human voluntary drug consumption^30^. Finally, we conducted whole-brain functional connectivity (FC) analyses employing complementary methods to assess both local and global changes in brain dynamics. In summary, we developed an NHP model of chronic cocaine use, abstinence and re-exposure that allows for whole brain interrogation of disease progression over a long timescale.

## Results

Four naive macaques (Macaca fascicularis: two female, and Macaca mulatta, two male) self-administered cocaine following an Intermittent Access protocol (IAp). The protocol consisted of defined intervals of access, i.e., 5 minutes of drug availability (or 20 lever presses; LPs), followed by 25 minutes of drug unavailability. Subjects were exposed to a series of six intervals of access and subsequent drug unavailability periods. The duration of every session was between 150 to 180 minutes (see **Fig. 1A** and Methods). Subjects could voluntarily trigger drug delivery via a reward box with a response lever, drug availability (red) and drug delivery (yellow) lights (**Fig. 1B** and Methods). The cocaine was delivered intravenously through a vascular access port (VAP, **Fig. 1C** and Methods). Each LP triggered the delivery of a cocaine infusion of 0.075 mg/kg. The maximum daily dose of cocaine was 9 mg/kg. The subjects were trained on the IAp with juice reward before the start of the cocaine SA (see **Fig. 1D** and Methods). We acquired rs-fMRI data while the subjects were lightly anesthetized (see **Fig 1D**. and Methods): 2 baseline scans before cocaine SA (at least 3 months apart); 3 scans during the SA period (i.e., after 5 days, 45 days and 95 days of cocaine use); 3 scans during the abstinence period (i.e., 5, 30 and 60 days after the end of the cocaine SA); 1 scan after re-exposure (i.e., 5 days of cocaine SA).

**Figure 1.**
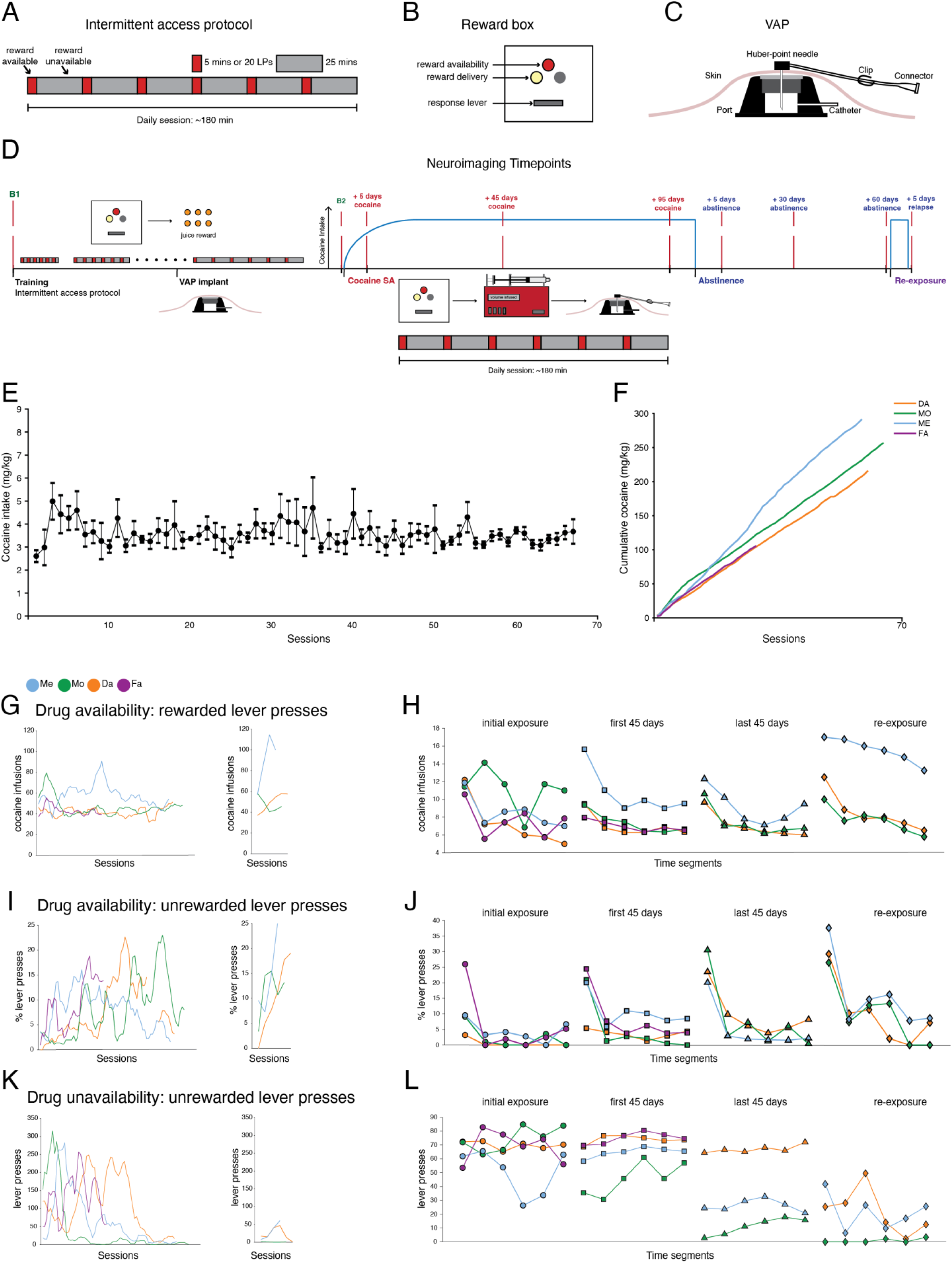
O*v*erview *of study design and self-administration behavior* **A** Intermittent access protocol for cocaine self-administration. The protocol included defined intervals of access, i.e., 5 minutes of drug availability, or 20 lever presses (LPs), each followed by 25 minutes of drug unavailability. Subjects were exposed to a series of six intervals of access and subsequent drug unavailability periods. The duration of every session was between 150 to 180 minutes. **B** Reward box for triggering cocaine delivery. Subjects were placed in an environment where they could voluntarily trigger drug delivery. This environment includes a reward box with a response lever, drug availability (red) and drug delivery (yellow) lights. Each LP triggered the delivery of a cocaine infusion of 0.075 mg/kg. **C** Vascular access port (VAP) for cocaine delivery. The VAP consists of an implanted reservoir (the port) under the skin that is connected to a catheter. The catheter leads into a large vein, allowing for direct access to the bloodstream. The VAP is accessed using a Huber point needle which has a unique, non-coring tip that is specifically designed to penetrate the silicone membrane of VAPs without causing damage to the port itself. **D** Training and neuroimaging timeline. The subjects were trained on the intermittent access protocol with juice reward before the start of the cocaine SA; the reward unavailability was progressively increased up to 25 mins based on performance (see **Methods**). The VAP was implanted between 6-8 weeks before the start of the cocaine SA. The first baseline scan (B1) was acquired at least 3 months before the start of the cocaine SA. The second baseline scan (B2) was acquired right before the start of the cocaine SA. The SA neuroimaging timepoints were after 5 days, 45 days and 95 days of cocaine. The abstinence timepoints were 5, 30 and 60 days after the end of the cocaine SA. The re-exposure consisted of five days of cocaine SA, followed by an additional scan. **E** Average daily cocaine intake over sessions. Cocaine maintained robust SA with no clear signs of escalation of use. Circles: average cocaine intake ± s.e.m across subjects. Y-axis: cocaine intake (mg/kg). X-axis: cocaine SA sessions. **F** Individual differences in cumulative cocaine intake over sessions. Lines (and colors): individual subjects, i.e., Da Mo, Me, Fa. Y-axis: cumulative cocaine intake (mg/kg). X-axis: cocaine SA sessions. **G** Cocaine SA over time during the initial self-administration and subsequent re-exposure phases. Y-axis: The session-total number of cocaine infusions. X-axis: Session number. Note: The time series were smoothed using a moving window of 5 sessions for the initial, and 2 sessions for the re-exposure phases. **H** The average within-session pattern of cocaine SA across time segments. Each line consists of the average cocaine SA across sessions, for each of the 6 blocks. Y-axis: The average number of cocaine infusions over blocks, across sessions. X-axis: Time segment. **I** Unrewarded lever presses during the drug availability period over time during the initial SA and subsequent re-exposure phases. The unrewarded lever presses were defined as those occurring during drug infusion. Y-axis: The percentage of unrewarded lever presses, i.e., unrewarded LPs/total LPs during the drug availability period. X-axis: Session number. Note: The time series were smoothed using a moving window of 5 sessions for the initial, and 2 sessions for the re-exposure phases. **J** The average within-session pattern of unrewarded LPs during the drug availability period. Each line consists of the average percentage of unrewarded LPs, for each of the 6 blocks. Y-axis: The average percentage of unrewarded LPs during the drug availability period. X-axis: time segment. **K** Unrewarded LPs during the drug unavailability period over time during the initial SA and subsequent re-exposure phases. Y-axis: The session-total number of unrewarded LPs during the drug unavailability period. X-axis: Session number. Note: The time series were smoothed using a moving window of 5 sessions for the initial, and 2 sessions for the re-exposure phases. **L** The average within-session pattern of unrewarded LPs during the drug availability period. Each line consists of the average unrewarded LPs, for each of the 6 blocks. The symbols represent the time segments. Circle: Initial 5 days of cocaine exposure; Square: The first 50 days of SA; Triangle: the last 50 days of self-administration; Diamond: the 5 days of re-exposure. Y-axis: The average unrewarded LPs during the drug unavailability period. X-axis: Time segment.

### Behavior

Cocaine maintained robust SA in all subjects (see **Fig. 1E**). In the first ∼5 sessions, the average cocaine consumption increased from ∼ 2.6 mg/kg to ∼ 5 mg/kg and subsequently stabilized to an average consumption of ∼4 mg/kg for the rest of the SA period. The average cocaine consumption across subjects and sessions was 3.54 ± 0.054 mg/kg. It is important to note that we did not observe an escalation of use which aligns with previous NHP cocaine SA studies^31,32^. By examining individual trajectories of cumulative cocaine intake, we observed subtle differences in the rate of cocaine consumption that developed over time, which were not apparent during the initial use phase (see **Fig. 1F**).

During chronic cocaine use, the behaviors of interest were: (1) rewarded LPs during drug availability across (**Fig. 1G**) and within sessions (**Fig. 1H**) (2) unrewarded LPs during drug availability across (**Fig. 1I**) and within (**Fig. 1J**) sessions; (3) unrewarded LPs during drug unavailability across (**Fig. 1K**) and within (**Fig. 1L**) sessions. The average cocaine consumption across sessions was stable across subjects except for Monkey Me who showed an increase in cocaine consumption around 45 days. This subject experienced technical difficulties with the VAP around that time, which may account for this observation.

Most importantly, the SA pattern observed within all sessions supports our IAp model as a more ecologically valid model of human cocaine consumption patterns (see **Fig. 1H**). In particular, the timing of the IAp was determined based on the peak cocaine plasma level and the corresponding drug-induced behavioral changes with intravenous cocaine administration in macaques (i.e., 2-5 mins after intravenous administration; hence the drug availability period was set to 5 mins), as well as the duration required for these effects to resolve (i.e., 15-30 mins)^33–35^. For all subjects, the highest level of cocaine SA occurred in the first availability interval and progressively decreased toward the end of the session. This pattern reflects both the spikes and falls, as well as the slow accumulation of plasma cocaine throughout the session, which accounts for the slow decrease in consumption ^34^.

Next, we examined the two behaviors related to unrewarded LPs which exhibited different timescales and patterns over time. During the availability period, unrewarded LP frequency was low for all subjects in the beginning (see **Fig. 1I)**. With prolonged use, this behavior exhibited oscillations and peaks at different times for each subject, with an overall increase in occurrence. Moreover, while each subject displayed a relatively similar pattern, the timing of the peaks was subject-dependent. Interestingly, the within-session pattern (see **Fig. 1J**) mirrored that of the cocaine infusions. This similarity suggests that unrewarded LPs during the drug availability might reflect compulsive drug-seeking behavior, or impaired response inhibition, that escalates with prolonged use. During the unavailability period, unrewarded LP frequency was high in the beginning for all subjects (see **Fig 1K**). With prolonged use, this behavior diminished and was nearly absent after 95 days of chronic cocaine use. Yet again, the behavioral trajectory and associated timescales were subject-dependent. During the re-exposure, this behavior remained low. The reduction in unrewarded LPs during the drug unavailability period might reflect the well-known transition between goal-directed and habitual drug use. Initially, despite knowing that no reward can be obtained in that interval, the motivating effects of cocaine might increase this behavior as a sign of frustration. With prolonged use and the shift towards habitual use, this behavior might diminish due to a loss of the motivational aspect of cocaine. There was no within-session pattern for this behavior.

### Local and global network changes relative to pre-drug baseline during chronic cocaine use, abstinence and re-exposure

To gain a comprehensive understanding of the changes induced by chronic cocaine use, abstinence and re-exposure, we used graph theory (GT) measures and FC magnitude changes (see **Methods** and **Fig. 2**). To provide a concise overview of the neurobiological changes associated with our experimental manipulations, we report our results with respect to the following functional networks: reward network (RN), habit network (HN), salience network (SAL), default mode network (DMN), central executive network (CEN), dorsal attention network (DAN), somatomotor network (SMN), multimodal integration network (MIN), auditory network (AUD) and visual network (VIS) ^10,21,36^. It is important to note that we did not derive these networks ourselves nor analyze our data using them as the unit of analysis but rather employed them as a framework to report our results (see **Supplementary Table 1**).

**Figure 2.**
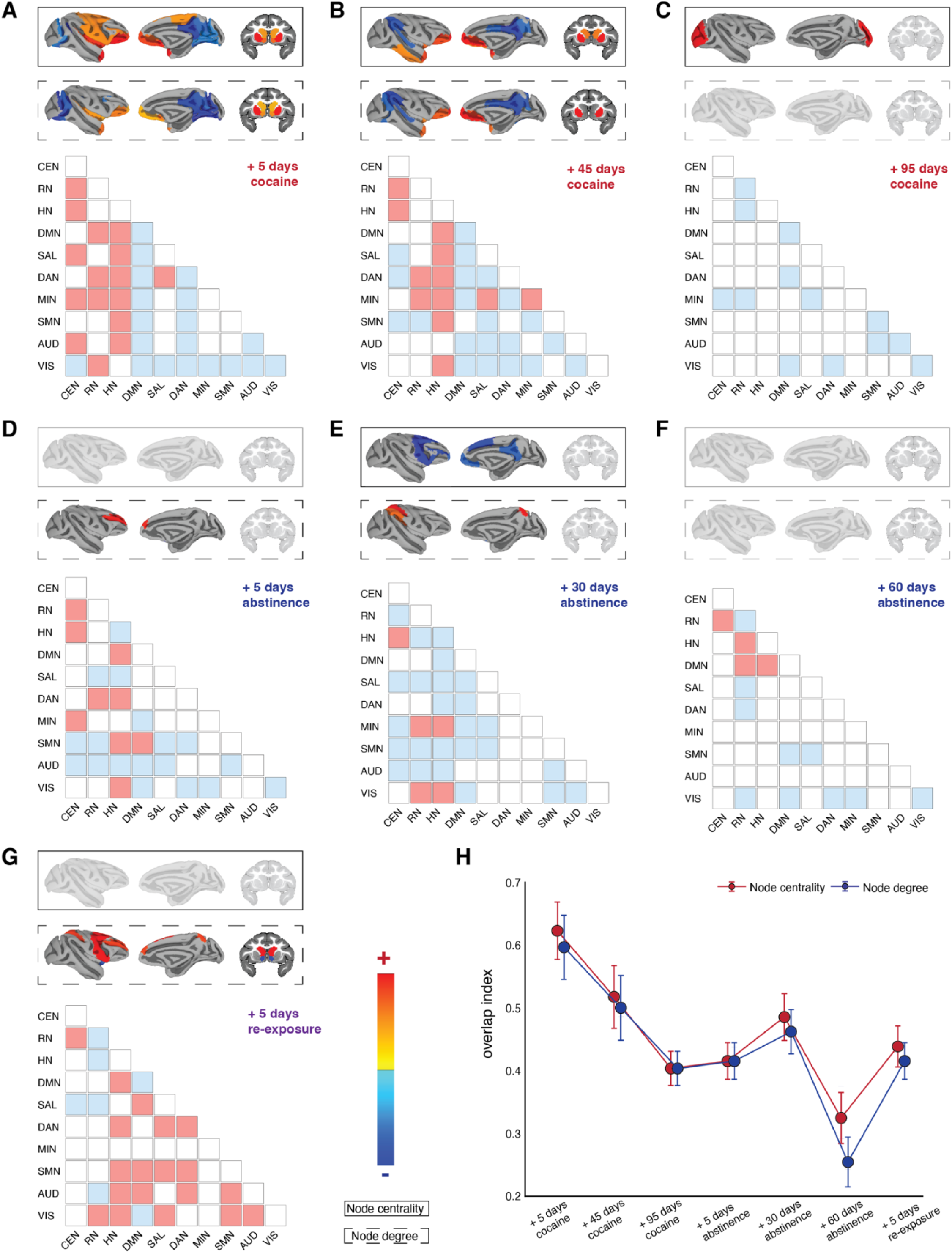
F*u*nctional *connectivity changes relative to pre-drug baseline during chronic cocaine use, abstinence and re-exposure* **Top:** Network topology (i.e., node centrality and degree) changes from baseline. Chronic cocaine use, abstinence and re-exposure induced shifts in connectivity patterns and network organization both locally (degree) and globally (centrality). At each time point, network measures were computed for the positive adjacency matrix. For each graph theory measure and time point, a two-sided paired t-test was used to examine the group-level changes from baseline. Results were thresholded using a familywise corrected p-FDR < 0.05. **Bottom:** Local functional connectivity magnitude changes from baseline within and between networks (see **Supplementary Fig. 1**A for the detailed connection-level changes and statistics). Results were thresholded using p < 0.05 connection-level threshold and a familywise corrected p-FDR < 0.05 cluster level threshold. Red: increase in FC. Blue: decrease in FC **A** 5 days of cocaine use. **B** 45 days of cocaine use. **C** 95 days of cocaine use. **D** 5 days of abstinence. **E** 30 days of abstinence. **F** 60 days of abstinence. **G** 5 days of re-exposure. **H** Overlap index for node centrality and degree across subjects. For each subject, time point and ROI, changes from baseline were coded as +1 for positive and-1 for negative effects. The overlap index was then calculated as the absolute value of the mean effect across subjects, ranging from 0, indicating no overall trend, to 1, indicating that all subjects exhibited either a positive or negative effect. Subsequently, the effect was averaged across ROIs. Y-axis: average overlap index across subjects and ROIs; X-axis: time points. Red: node centrality; Blue: node degree.

After 5 days of cocaine SA (see **Fig. 2A**), we found the following changes relative to baseline: (1) An increase in the FC of the CEN, RN and HN with the rest of the brain. (2) A reduction in DMN within-and across-networks FC. (3) A reduction in FC between sensory-motor and attention-related networks. (4) An increase in the node centrality of RN (areas 25, 11,13, 12, 14,10, 32, nucleus accumbens), HN (caudate, putamen), CEN (vlPFC), MIN (TGa, TGg/d), SAL (insula), and SMN (PM, S1, S2, M1) areas; (5) A decrease in node centrality for DMN (PCC, RSC, area 7), DAN (vmIPS), VIS (V2, V3, V6 V6A, MT, MST) and AUD (belt) areas; (6) An increase in node degree of RN (areas 11/13, 25, 12, 10, 14, nucleus accumbens), HN (putamen, caudate), MIN (TGg/d), and SAL (insula) areas. (7) A decrease in node degree for DMN (PCC, RSC, area 7), DAN (IPS, FEF), VIS (V2, V3, V4, V6, V6A, MT, MST) and AUD (belt) areas. Overall, we observed a homogenous effect following the initial use of cocaine, which was characterized by an increased role of RN, HN, CEN and SMN areas, and a decreased role of the DMN, DAN, VIS and AUD areas in network dynamics. The FC magnitude changes offer further insights into the observed shifts in GT measures. The increased role of RH, HN and CEN is mediated by enhanced FC with most brain regions, while the self-directed, attention, salience, sensory and somato-motor networks exhibit reduced connectivity with each other.

After 45 days of cocaine SA (see **Fig. 2B**), we found the following changes relative to baseline: (1) An increase in FC between CEN, RN, HN with the rest of the brain. However, this was less pronounced than following just 5 days of cocaine SA. In particular, we observed a shift toward decreased FC between CEN and the other networks. (2) Similar to initial cocaine use, the DMN, sensory, somato-motor and attention-related networks displayed reduced FC with each other. (3) An increase in the node centrality of RN (area 25, 32, 14, 10, 12, 11, 13), HN (caudate, putamen), and MIN (TGd/g, TGa) areas. (4) A decrease in node centrality of DMN (PCC, RSC, area 7), DAN (IPS), SMN (MCC), SAL (IPL), and VIS (MT, MST, V6A, TE) areas. (5) An increase in the node degree of RN (area 25, 32, 11, 13, 10, 12), HN (putamen), and MIN (TGd/g, TGa) areas. (6) A decrease in the node degree of DMN (PCC, RSC, area 7), DAN (IPS), VIS (V4, MT, V6, V6A) and AUD (STG) areas. So far, our observations reveal a distinct separation in the effects of cocaine on brain activity; specifically, RN, CEN, and HN were hyper-engaged, while the DMN, sensorimotor, and attention networks were hypo-engaged. However, despite similarities across time points, prolonged cocaine exposure resulted in less pronounced effects.

After 95 days of cocaine SA (see **Fig. 2C**), we found only the following changes relative to baseline: (1) A decrease in FC across CEN, RN, HN, DMN, SAL, SMN, MIN, VIS and AUD. Overall, we observed a major attenuation of group-level significant results. Importantly, we observed a more localized connectivity decrease at the within-network level rather than across networks. (2) An increase in the node centrality of V1.

During early abstinence (see **Fig. 2D**), we found the following changes relative to baseline: (1) Increase in the FC between CEN and the RN, HN and MIN. (2) In contrast to drug use, a decrease in FC between CEN and sensory and somato-motor networks. (3) CEN hyperactivity with respect to RN, HN and MIN. (4) An increase in FC between HN and VIS, DMN, DAN, CEN and DMN. (5) A decrease in FC between RN and AUD, SMN, and SAL. (6) Decrease in DMN FC with the rest of the brain, except with HN and SMN. (7) A decrease in FC between VIS, AUD, SMN, DAN with the rest of the brain. (8) An increase in the node degree of dlPFC (CEN). In contrast to the previous timepoint, at the beginning of abstinence we observed widespread changes in FC with GT measures being impacted in only one area. Connection-level changes can occur without corresponding changes in GT measures due to the distinction between localized alterations and global network properties.

After 30 days of abstinence (see **Fig. 2E**), we found the following changes relative to baseline: (1) Overall decrease in FC across the whole brain. The only exceptions were RN and HN, which displayed an increased FC with MIN and VIS, and HN which displayed an increased FC with CEN. (2) a decrease in the node centrality of RN (area 10), CEN (vlPFC, dlPFC), DMN (PCC, RSC), and SMN (PM, SMA) areas. (3) An increase in the node degree of other SMN (area 5, retroinsula) and SAL (IPL) areas.

After 60 days of abstinence (see **Fig. 2F**), there were no results for the GT measures. With respect to FC magnitude changes (see **Fig. 2F**), the effects were notably attenuated. RN exhibited an increase in FC with HN, DMN and CEN but an overall within-network decrease. SMN and VIS displayed a decrease in FC.

Finally, after 5 days of re-exposure (see **Fig. 2G**), we found the following changes relative to baseline: (1) Increase in FC between the DMN, SAL, DAN and VIS, AUD, and SMN, which contrasted with initial cocaine use. (2) Increase in FC between HN and most networks. (3) Increase in FC between RN and CEN and a decrease in FC within RN itself. (3) An increase in the node degree of HN (caudate), CEN (dlPFC) and SMN (PM, area 5) areas. (4) A decrease in the node degree of RN (nucleus accumbens) areas.

### Heterogeneity of individual trajectories in functional connectivity across chronic cocaine use, abstinence and re-exposure

Thus far, we have shown that, with prolonged cocaine use, the initial group-level effects diminished. Likewise, the trajectory of abstinence-induced neural alterations was nonlinear but distinct to that observed during chronic cocaine use. Group-level effects only emerged after 1 month of abstinence, but by 2 months, these effects attenuated. We hypothesized that drug-induced, and subsequent abstinence-induced, alterations in brain function, might vary across time and subjects. This would manifest as increased heterogeneity of effects across subjects.

To that end, we computed an overlap index to assess how the overall direction of changes from baseline varies across subjects over time, without considering the magnitude of the effects (see **Fig. 2H**). We found that the heterogeneity of effects was lowest during initial cocaine use and progressively increased with prolonged use. Abstinence-induced neural alterations converged across subjects after one month, however by 2-months, individual trajectories reveal heterogeneity, suggesting differences in recovery. During re-exposure, heterogeneity decreased again. With continued use, the increase in the diversity of effects demonstrates the complexity of drug-induced neural alterations, where individuals’ neural trajectories begin to diverge potentially based on interindividual differences such as genetic and environmental factors. With respect to abstinence, the individuals may initially experience similar changes in brain function in the beginning of recovery. Nevertheless, longer-term recovery could be influenced more heavily by a variety of individual factors yet to be determined. The decrease in heterogeneity during the re-exposure phase suggested that individuals exhibit similar re-exposure-induced neural alterations.

Finally, we were interested in whether these heterogeneity results could be explained by reduced impact on the brain of drug (or abstinence), or whether brains were still responding and changing, but heterogeneously. We examined how node degree and centrality effects changed across time at the individual subject level (see **Supplementary Fig. 1B**). We did not observe a linear diminishing effect in any of the subjects, but rather nonlinear dynamics across the stages of substance use, abstinence and re-exposure. We thus concluded that subject’s brains were still responding to drug/abstinence but were doing so in different ways.

### Linking heterogeneity in brain changes to behavioral variability during chronic cocaine use

Next, we investigated whether the heterogeneity of neural responses during drug use was related to behavioral variability across subjects (see **Methods** and **Fig. 3**). To assess whether individual differences in behavior explain the variability in FC changes from baseline, we looked at the node degree change from baseline (see **Methods** and **Fig. 2**). This was done to obtain a directionally unbiased measurement of how behavior could be related to network changes agnostic of the region’s place within a network. The behavior was divided into 4 segments: before the first neuroimaging time point, SA between the first and second neuroimaging time points, SA after the second neuroimaging time point, and after the re-exposure time point. For each segment and NHP, the behavioral variables were summed across intervals and averaged across sessions. For each timepoint and region-of-interest (ROI), a multiple regression model was fitted with the three behavioral measures as predictors and changes in node degree as the dependent variables. Permutation testing was conducted to assess significance, and a secondary threshold criterion was applied, involving the selection of models within the top 25th percentile across ROIs (see **Fig. 3**).

**Figure 3.**
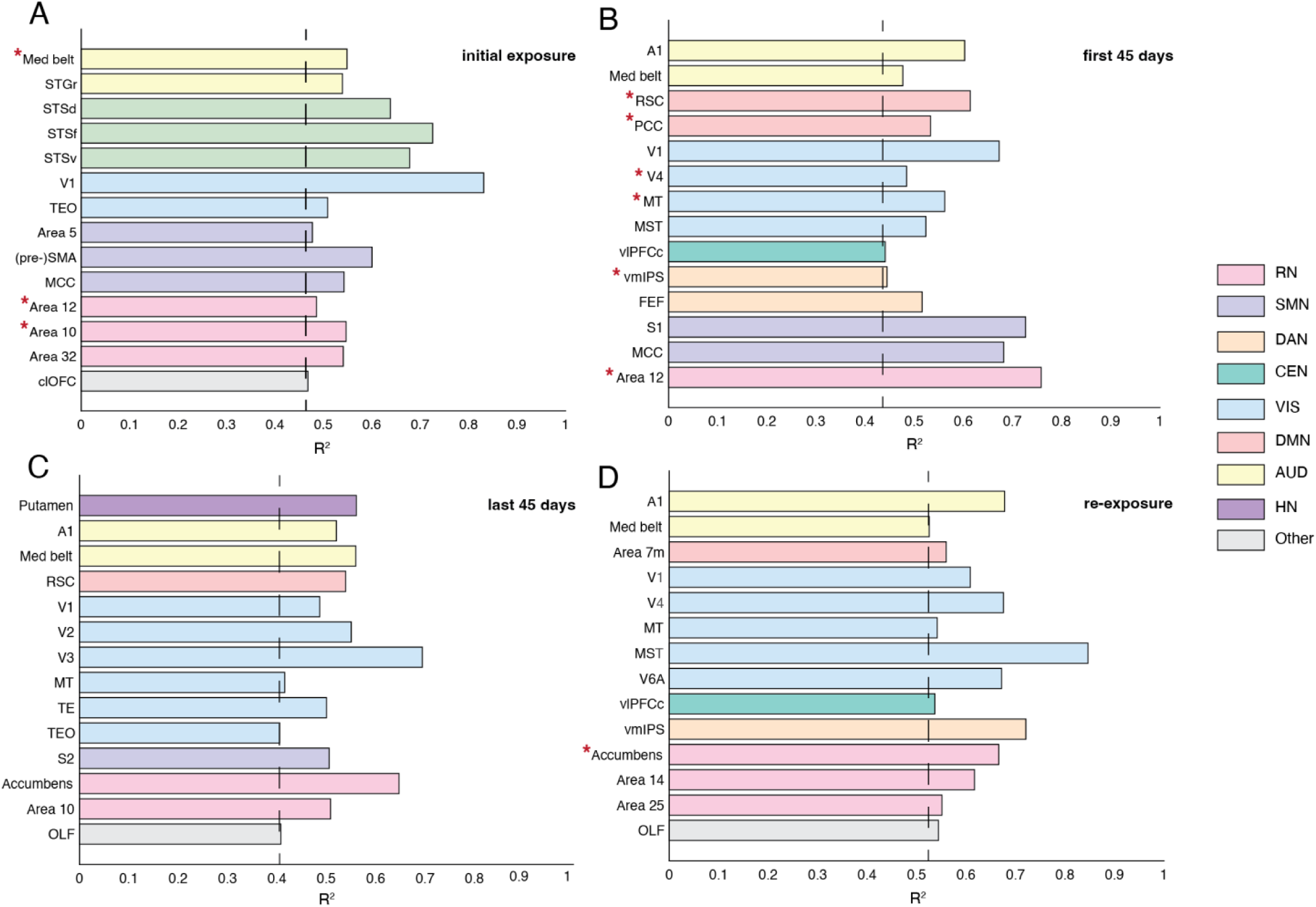
I*n*dividual *differences in chronic cocaine self-administration behavior and neural changes* Individual differences explain the variability in degree changes from baseline. The behavioral measures were the unrewarded and rewarded LPs during the drug availability period and the TO LPs. The behavior was divided into 4 segments: SA before T1 (A), SA between T1 and T2 (B), SA after T2 until the end (C), and re-exposure (D). For each ROI, the resulting behavioral variables were used as predictors in a multiple regression model with the change in node degree as dependent variable. We used permutation testing to assess the statistical significance of the observed relationships between behavioral predictors and changes in graph theoretical measures (see **Methods**). Furthermore, we established a secondary threshold criterion across ROIs, whereby only ROIs exhibiting an R^2 value within the top 25th percentile of ROI models was deemed robust and indicative of significant predictive power. In each panel, the dotted line represents the 75^th^ percentile across ROIs. Y-axis: ROIs. X-axis: R^2 of the multiple regression model. Areas exhibiting both significant brain-behavior associations and group-level changes from baseline in degree are marked with an asterisk (*).

After 5 days of cocaine SA (see **Fig. 3A**), the 75^th^ percentile threshold was R^2 = 0.47. The behavior accounted for variability in individual brain responses to cocaine use in the RN, SMN, MIN, VIS and AUD networks. We have shown that areas in these networks exhibit homogenous changes in node degree from baseline across subjects (see **Fig. 2A**). However, most areas with a brain-behavior association did not show significant group-level changes in node degree – i.e., 11 out 14 areas. For areas such as 10 and 12, we observed significant changes in both node degree from baseline and brain-behavior associations. This suggests that some areas may exhibit consistent cocaine-induced changes that are nonetheless graded across subjects. In other words, while the effects of cocaine on certain ROIs were similar across subjects, the magnitude of that effect varies across individuals. Interestingly, the striatum did not show an association with behavioral heterogeneity; instead, it was strongly homogenous across individuals. Overall, after initial exposure to cocaine, the behavior was predictive of changes in node degree in reward, sensory and motor areas.

After 45 days of cocaine SA (see **Fig. 3B**), the 75^th^ percentile threshold was R^2 = 0.44. Behavior accounted for variability in individual brain responses to cocaine use in the RN, CEN, SMN, DAN, DAN, VIS and AUD networks. We observed a similar pattern as in the 5-day timepoint, where most areas that show significant associations to behavior do not exhibit significant group-level results – i.e., 8 out of 14. Notably, with prolonged cocaine use, behavior became more predictive of node degree changes in higher-order association areas underlying attention, self-regulation and decision-making. Unimodal sensory-motor areas remain closely linked to behavior.

After 95 days of cocaine SA (see **Fig. 3C**), the 75^th^ percentile threshold was R^2 = 0.41. Behavior accounted for variability in individual brain responses to cocaine use in the RN, HN, SMN, DMN, VIS and AUD.

After 5 days of re-exposure (see **Fig. 3D**), the 75^th^ percentile threshold was R^2 = 0.52. The behavior accounted for variability in individual brain responses to cocaine use in the RN, CEN, DMN, VIS and AUD networks. We observed the same pattern wherein none of the areas that show a significant association with behavior display group-level significant results.

Overall, these results underscore the limited overlap between group-level results and brain-behavior associations, suggesting that while a broader range of brain areas may be affected, only the commonalities at the group level are observed.

## Discussion

We established a preclinical model of chronic cocaine use that differentiates itself from existing research in three significant ways, mitigating limitations present in both human and current animal studies. First, we employed drug-naïve NHPs, which allows for the examination of cocaine-induced effects without the confounding influence of prior drug exposure. This is complemented by the fact that all subjects share similar use characteristics, including duration and pattern of cocaine use and the number of abstinence and re-exposure episodes. Such variables are challenging to address in human research and typically limit SUD findings ^37^. Second, we used a within-subject longitudinal design with subject-specific baselines instead of using a control group for comparison, enhancing the sensitivity of our findings. Matching user populations to controls is extremely challenging, especially in the context of SUDs, which can introduce confounding variables that may compromise the robustness of statistical analyses. Finally, we established a longitudinal model of chronic cocaine use, abstinence and re-exposure over a long timescale, combined with dense temporal sampling. This approach allows for a comprehensive examination of the dynamic changes associated with addiction phenomenology, offering unprecedented insights into the neural and behavioral progression that comes with chronic cocaine use.

Using our model, we found consistent brain-wide impairments in function, suggesting broad multi-faceted drug-induced alterations. However, while initial cocaine use resulted in convergence of effects across subjects; with prolonged use the brain responses to cocaine SA diverged. One implication of our findings is that heterogeneity in SUD populations may not solely stem from preexisting individual differences, but also from diverse neural responses to drug exposure. In contrast, early abstinence was characterized by divergent brain responses across subjects, while after one month of abstinence, we observed a convergence of the effects across subjects. After 2 months of abstinence, we found divergent individual trajectories that could suggest differences in the timescales and extent of recovery.

Recent research highlighting the significant heterogeneity characteristic of psychiatric disorders suggests that group-level analyses using traditional case-control paradigms with large sample sizes may be a key factor contributing to the limited progress in the field (Segal et al., 2025). While ethically and methodologically challenging in humans, baseline assessments in animal studies are crucial for understanding the neurobiological changes associated with any experimental manipulation. It is therefore essential to move beyond the notion that interindividual differences are merely noise and embrace them in psychiatric research ^24,38,39^. To enhance the translational value of our preclinical model of chronic cocaine use, we addressed the significant heterogeneity in SUDs by employing dense neuroimaging sampling coupled with long sessions and high SNR afforded by ultra-high magnetic field strengths ^21,23,40^. We focused our analyses on local and global whole-brain network dynamic changes from the individual-specific baselines. We posit that to identify MRI biomarkers related to any psychiatric disorder, we must leverage both group commonalities and the heterogeneity in individual-specific features. Hence, we investigated both group-level and individual trajectories throughout chronic cocaine use, abstinence and re-exposure.

We observed a strong homogenous effect across subjects in response to initial use of cocaine, however the individual trajectories diverged with prolonged use and group-level results diminished as a result. Our findings are consistent with a recent review in rodents which revealed that reward areas dynamically adapted to chronic drug use depending on the length of administration ^41^. In the striatum, rodent studies with cocaine ^42^ and human studies of stimulant administration ^43^ revealed differential effects following acute versus chronic stimulant use. We extend those results by demonstrating that the whole brain dynamically adapts to chronic cocaine use over a long timescale. However, this adaptation does not include a return to baseline. Instead, each individual brain responded uniquely to chronic cocaine use. The divergence of individual neural trajectories has crucial implications for developing treatments since what might work in short-term users might not work in long-term users ^43^.

We demonstrated that abstinence exhibited a similarly nonlinear and nonstationary effect. Abstinence-induced neural alterations converged across subjects after 1-month, but by 2 months, individual trajectories were heterogenous, possibly indicating differences in recovery. We posit that the early abstinence results could be related to interindividual differences in incubation of craving which has been shown to be susceptible to variation in magnitude as a function of biological or environmental circumstances surrounding the individual ^44^. These observations raise the question of whether dynamic adaptation occurs on different timescales for individual subjects or if the changes are simply different across subjects.

We bring evidence that dynamic neural and behavioral adaptations occur on different timescales for individual subjects. We found that most areas exhibiting significant changes in degree from baseline at the group-level, do not demonstrate significant brain-behavior associations across subjects. This suggests that individual-specific effects linked to behavior may be less detectable at the group-level, emphasizing the need for analyses that preserve and investigate individual variability. This observation, combined with the attenuation of group-level effects over prolonged cocaine use and the varying individual behavioral timescales, supports the conclusion that cocaine-induced effects become more individualized over time. Group analyses could mask these nuances. This underscores the importance of individualized approaches in studying SUD by considering the unique trajectories and timescales of neural and behavioral changes in each subject. Importantly, for a small number of ROIs, we observed both significant changes in node degree from baseline and brain-behavior associations. This suggests that certain ROIs demonstrated graded effects. Essentially, these areas exhibited effects in the same direction that varied in magnitude across subjects. However, this pattern is only present during initial exposure; and after 45 days. Furthermore, certain areas transition from exhibiting graded responses with consistent directionality across subjects to displaying brain-behavior associations with divergent directions, ultimately losing group-level significance. This progression provides additional evidence that neural response heterogeneity increases over time, shifting from shared, graded effects to individualized, divergent patterns that diminish group effects. Lastly, this phenomenon is not limited to a specific network but is observed across the whole brain, indicating a widespread increase in neural response heterogeneity with prolonged exposure. For example, initially, the striatum reflects strong, uniform effects of cocaine across subjects. However, with prolonged use, heterogeneity emerges within the putamen, and this heterogeneity is linked to behavioral variability. Interestingly, at this timepoint, our results align with previous findings demonstrating an association between cocaine consumption and changes in FC in the putamen ^32^. Following re-exposure, a re-establishment of homogeneity occurs in the caudate nucleus, accompanied by brain-behavior associations in the nucleus accumbens, suggesting a dynamic reorganization of striatal circuits in relation to addiction progression. It is important to emphasize that what we observe with respect to interindividual differences in behavior is not fundamentally a different pattern but rather different timescales over which the behaviors develop. Yet again, we demonstrate the importance of considering the temporal component in psychiatric analyses ^45^. In sum, we highlight the limited overlap between group-level results and brain-behavior associations, indicating that only common effects are captured at the group-level despite widespread brain involvement.

We found that prolonged use of cocaine shifts the brain towards a widespread desynchronization. This finding has also recently been reported with recreational use of psychedelics in humans ^14,67–7046^. Our results however highlight the need for caution in generalizing the effects of a drug based on occasional use, as such assumptions may not accurately reflect the broader implications of prolonged use. We show that initial use of cocaine synchronizes reward and higher order prefrontal areas and desynchronizes sensory and self-related areas, suggesting an imbalance between reward and executive networks and self-referential and sensory areas. However, prolonged use of cocaine shifts the effects towards a widespread desynchronization of the brain. The re-exposure dynamics suggest a noteworthy connection to initial cocaine use, in that both induce a strong homogenous effect across subjects, despite the significant differences in the underlying neural and behavioral effects associated with them. This link suggests that while the specifics of how chronic drug use and its consequences may significantly vary across individuals, the immediate, acute effects of cocaine have a consistent impact when used for the first time or after an abstinence period. This finding highlights that cocaine’s pharmacological effects can only override individual differences with initial use or after a period of abstinence. Understanding these dynamics is critical for developing effective interventions that consider both the homogeneity of drug effects and the individuality of drug use and recovery trajectories. A critical piece of information is that initial use and re-exposure exhibited almost opposite effects on the brain. That implies that drug history and polydrug use is a larger con^14,14,67–70found^ than we previously thought.

In conclusion, we set out to map the addiction connectome; however, the results indicate that the neural correlates of SUD are more complex than can be captured by a singular, canonical connectome. It is thus more accurate to state that we mapped the progression of the macaque functional connectome from a naive state through chronic cocaine use, abstinence and re-exposure, providing a comprehensive characterization of each phase. Our primary finding is that the addiction connectome is a dynamic interaction of subject by drug by time; the effects vary and evolve dynamically throughout the stages of chronic cocaine use and recovery. Second, we observe that the convergence of effects across subjects also shifts dynamically over time, indicating that the magnitude and direction of cocaine’s impact vary not only within the group but also across subjects. This distinguishes from the first point by emphasizing that the effects are not only non-specific to particular regions but also exhibit temporal variability in their consistency and direction across different subjects. Third, we find that the divergence across subjects is partly explained by heterogeneity in behavior; however, it is important to note that all subjects exhibit a similar progression of behavioral changes, though these occur over different timescales. This distinction is crucial because it suggests that while individual differences influence the timing of behavior, the behavioral trajectory is consistent across subjects. This indicates that there are two distinct types of individual differences: first, variability in the specific effects of cocaine on neural and behavioral measures (heterogeneity in cocaine-induced effects), and second, differences in the timing or rate at which these effects manifest (heterogeneity in the timescales of cocaine-induced effects). While our data suggest the presence of these two sources of variation, further research is necessary to fully understand their implications. To address this, future research should focus on comprehensive behavioral and neural phenotyping, incorporating a broader array of measures that assess multiple cognitive domains. Enhanced sampling density is essential for accurately capturing the neural and behavioral trajectories over time, while simultaneously monitoring heterogeneity in effects across subjects. This approach will facilitate the disentanglement of effects that are fundamentally similar across individuals but manifest over varying timescales, from those that are consistently different between subjects. Finally, we emphasize the clinical promise of precision functional mapping that is performed over long timescales to help guide personalized medicine and provide a higher standard of care. It is imperative to differentiate between identifying targets for interventions that are common across subjects, and developing interventions tailored to the individual and their associated timescales. We and others ^24,38,45^ propose that the current case-control paradigm is inadequate to capture the inherent biological and clinical variability of psychiatric disorders. We argue that even with an increased sample size to increase statistical power, group-level analyses may still fall short compared to precision mapping techniques. While increasing the sample size appears to increase statistical power for finding extremely small effects ^47^, it also obscures meaningful interindividual differences, reducing them to mere noise. Hence, brain wide association studies are unlikely to provide the nuanced insights necessary for our understanding of psychiatric disorders, and subsequent identification of targets of intervention. Finally, our findings complement those emerging about neuroimaging subtyping in psychiatric disorders ^48^. The potential of neuroimaging-based subtyping lies in identifying groups of individuals sharing common biological signatures, thereby facilitating the development of biologically-informed, targeted treatments. Subtyping occupies an intermediate position between precision neuroimaging and brain-wide association studies serving as a compromise for developing targeted interventions that are as individualized as possible. Nonetheless, animal research is essential for informing these studies and testing hypotheses, as it provides a means to establish causality.

## Materials and Methods

Experimental procedures were carried out in accordance with the University of Minnesota Institutional Animal Care and Use Committee and the National Institute of Health standards for the care and use of NHPs. All subjects were fed ad libitum and single-housed within a light and temperature-controlled colony room. Animals had access to ad lib water.

### Subjects

We collected data from 4 macaque monkeys (Macaca fascicularis: two female, and Macaca mulatta, two male) without any prior nonclinical drug exposure, implant or cranial surgery, or behavioral training. Each monkey was fitted with an aluminum collar and trained to approach the front of the cage when the investigator was present. A stainless-steel rod with a latch on the end was attached to the collar and the NHP was guided into the primate restraint chair. The NHPs were placed in an environment where they could voluntarily trigger reward delivery. This environment included a reward box with a response lever, reward availability (red) and reward delivery (yellow) lights (see **Fig. 1B**). LP triggered the delivery of reward.

### Drugs

Cocaine hydrochloride was in 0.9% saline solution with a concentration of 10 mg/ml. Cocaine doses were determined based on the free base weight.

### Self-administration protocol

We used an IAp for cocaine SA (see **Fig. 1A**). The protocol included defined intervals of access, i.e., 5 minutes of drug availability, or 20 LPs, followed by 25 minutes of drug unavailability. Subjects were exposed to a series of six intervals of access and subsequent drug unavailability periods. The duration of every session was between 150 to 180 minutes. The maximum dose of cocaine that could be obtained in a session was 9 mg/kg (0.075 mg/kg per infusion).

#### Training

The NHPs were initially trained to respond to the lever when the reward light was on, by reinforcing each response with a set amount of preferred juice. When the NHPs exhibited consistent lever pressing, we started training them on the IntA protocol. The NHPs were initially trained on a 5-min reward availability and 5-min reward unavailability. As previously mentioned, subjects were exposed to a series of six intervals of access and subsequent reward unavailability periods. The duration of the reward unavailability period was gradually increased to 25-min based on performance. Notably, the reward unavailability period was increased when > 80% of the total LPs in a session were in the juice availability period. Training was completed when the NHPs would reach the above-mentioned performance with a 5-min reward availability and 25-min reward unavailability. During the experimental sessions, each LP triggered the delivery of a cocaine infusion of 0.075 mg/kg.

### Surgery and vascular access port

Each monkey was surgically implanted with a VAP (Access Technologies) for cocaine delivery. The VAP consisted of an implanted reservoir under the skin (i.e., the port) that was connected to an intravenous catheter. The catheter led into a large vein (i.e., saphenous vein distal to the knee), allowing for direct access to the bloodstream (see **Fig. 1C**). The VAP was accessed using a right-angle Huber Point Needle (Access Technologies) which has a unique, non-coring tip that is specifically designed to penetrate the silicone membrane of VAPs without causing damage to the port itself. Prior to accessing the port, the skin around the VAP was cleaned with 2% chlorhexidine solution before needle insertion. To ensure proper maintenance of the VAP during the cocaine SA sessions, the VAP was flushed with saline and filled with either TCS catheter lock solution (Taurolidine and Citrate; Access technologies) or Alteplase (Activase) daily to prevent clot formation and bacterial/fungal growth. To ensure proper maintenance of the VAP outside the cocaine SA sessions, the port was flushed with saline and filled with Alteplase monthly.

### Study design

Experimental sessions were conducted on average 5 days/week. Over a nine month period, anatomical and functional MRI scans were acquired at 9 time points: 2 baseline scans before initial drug use (at least 90 days apart); 1 after 5 days of drug SA; 1 after 45 days of drug SA; 1 after 95 days of SA; 1 after 5 days of abstinence; 1 after 30 days of abstinence; 1 after 60 days of abstinence and 1 scan after re-exposure (see **Fig. 1D**). The re-exposure phase consisted of 5 SA sessions and took place immediately after the 8th neuroimaging timepoint. For the re-exposure phase, the NHPs were placed back in the environment associated with the initial cocaine use and underwent the same IntA SA protocol. Every scan was conducted after 2 days of cocaine unavailability to avoid the effects associated with acute abstinence—i.e., the “crash” (1-2 days) which is characterized by an immediate post-cocaine depression following a binge, with symptoms such as drug craving, depression, agitation, and anxiety. During the abstinence phase, the NHPs were not brought back to the experimental room and only left their home cage for the MRI scans.

### MRI data acquisition

On scanning days, anesthesia was first induced by intramuscular injection of atropine (0.5 mg/kg), ketamine hydrochloride (7.5 mg/kg), and dexmedetomidine (13 µg/kg). Initial anesthesia was maintained using 1.0%–2% isoflurane mixed with oxygen. For functional imaging, the isoflurane level was lowered to 1%. All data were acquired on a passively shielded 10.5 T, 88-cm diameter clear bore magnet coupled to Siemens gradients (‘SC72’ body gradients operating at a slew rate of 200 mT/m/s, and 70 mT/m maximal strength), and electronics (Magnetom 10.5 T Plus) (Siemens, Erlangen, Germany). Within the gradient set and the bore-liner, the space available for subject insertion had a 60-cm diameter.

The 10.5-T system operates on the E-line (E12U) platform which is directly comparable to clinical platforms (3 T Prisma/Skyra, 7 T Terra). As such, the user interface and pulse sequences were identical to those running on clinical platforms. A custom in-house built and designed RF coil with an 8-channel transmit/receive end-loaded dipole array of 18-cm length (individually) combined with a close-fitting 16-channel loop receive array head cap, and an 8-channel loop receive array of 50×100 mm2 size located under the chin (Lagore et al., 2021). The size of 14 individual receive loops of the head cap was 37 mm with two larger ear loops of 80 mm—all receiver loops were arranged in an overlapping configuration for nearest neighbor decoupling. The resulting combined 32 receive channels were used for all experiments and supported threefold acceleration in phase encoding direction. The coil holder was designed to be a semi-stereotaxic instrument holding the head of the animal in a centered sphinx position via customized ear bars. The receive elements were modeled to adhere as close to the surface of the animal’s skulls as possible. Transmit phases for the individual transmit channels were fine-tuned for excitation uniformity for one representative mid-sized animal and the calculated phases were then used for all subsequent acquisitions. Magnetic field homogenization (B0 shimming) was performed using a customized field of view with the Siemens internal 3D mapping routines. Multiple iterations of the shims (using the adjusted FOV shim parameters) were performed, and further fine adjustment was performed manually on each animal. Third-order shim elements were ignored for these procedures.

In all animals, a B1+ (transmit B1) fieldmap was acquired using a vendor-provided flip angle mapping sequence and then power calibrated for each individual. Following B1+ transmit calibration, 3–5 averages (23 min) of a T1 weighted magnetization prepared rapid acquisition gradient echo protocol (3D MP-RAGE) were acquired for anatomical processing (TR=3300 ms, TE=3.56 ms, TI=1140 ms, flip angle=5°, slices=256, matrix=320×260, acquisition voxel size=0.5×0.5×0.5 mm3). Images were acquired using in-plane acceleration GRAPPA=2. A resolution and FOV matched T2 weighted 3D turbo spin-echo sequence (variable flip angle) was run to facilitate B1 inhomogeneity correction.

Before the start of the functional data acquisition, five images were acquired in both phase-encoding directions (R/L and L/R) for offline EPI distortion correction. Six runs of fMRI time series, each consisting of 700 continuous 2D multiband (MB)/Simultaneous Multi SLice (SMS) EPI (Moeller et al., 2010; Setsompop et al., 2012; Uğurbil et al., 2013) functional volumes (TR=1110 ms; TE=17.6 ms; flip angle=60°, slices=58, matrix=108×154; FOV=81×115.5 mm2; acquisition voxel size=0.75×0.75×0.75 mm3) were acquired. Images were acquired with a left-right phase encoding direction using in-plane acceleration factor GRAPPA=3, partial Fourier=7/8th, and MB or simultaneous multislice factor=2 (i.e., the number of simultaneously excited slices=2). Since macaques were scanned in sphinx positions, the orientations noted here are what is consistent with a (headfirst supine) typical human brain study (in terms of gradients) but translate differently to the actual macaque orientation.

In one subject (macaca mulatta), data collection was discontinued after the second cocaine time point (i.e., after 33 SA sessions) due to VAP complications. Additional neuroimaging data was acquired after 2 months of abstinence.

### MRI preprocessing

Image processing was performed using a custom pipeline relying on FSL^49^, ANTs^7450^, AFNI ^51^ and a heavily modified CONN^52^ toolbox. Images were denoised using Noise Reduction with Distribution Corrected (NORDIC) PCA ^53,54^. Images were then motion corrected using mcflirt (registration to the first image). Images were slice-time corrected and EPI distortion corrected using topup. High magnetic fields and large matrix sizes are commonly associated with severe EPI distortions. Anatomical images were nonlinearly warped into the ‘National Institute of Mental Health Macaque Template’ (NMTv2)^55^ template using ANTs and 3DQwarp in AFNI. The distortion correction, motion correction, and normalization were performed using a single sinc interpolation. Images were spatially smoothed (FWHM=2 mm), linear detrended, denoised using a linear regression approach including a five-component nuisance regressor of the masked white matter and cerebrospinal fluid and band-pass filtering (0.008–0.09 Hz)^56^.

### First-level analyses

ROI-to-ROI connectivity (RRC). RRC matrices were estimated characterizing the FC between each pair of regions among 57 ROIs (see Supplementary Table 1). FC strength was represented by Fisher-transformed bivariate correlation coefficients from a general linear model (weighted-GLM), estimated separately for each pair of ROIs, characterizing the association between their BOLD signal time series.

### Group-level analyses

To mitigate the small sample size, leverage the large amount of data we acquired for each individual, and account for within-session variability, we used the runs within each subject as independent observations, i.e., 24 observations for N = 4, and 18 observations for N = 3. Importantly, treating runs as separate subjects allows for correcting for within-session variability. Given within-subject variability across time, analyzing runs separately can increase the sensitivity of detecting significant cocaine-induced changes. Nevertheless, it is important to acknowledge that this approach may limit the generalizability of the results to a broader population. However, considering recent research demonstrating the substantial heterogeneity that characterizes psychiatric illness, group-level analyses with traditional case-control paradigms and a large N are potentially a major reason for the limited progress in the field^38^.

#### RRC

Group-level analyses were performed using a GLM (Nieto-Castanon, 2020). For each individual connection a separate GLM was estimated, with first-level connectivity measures at this connection as dependent variables (one independent sample per subject and one measurement per experimental condition), and timepoints as independent variables. Connection-level hypotheses were evaluated using multivariate parametric statistics with random-effects across subjects and sample covariance estimation across multiple measurements. Inferences were performed at the level of individual clusters (groups of similar connections). Cluster-level inferences were based on parametric statistics within-and between-each pair of networks (Jafri et al., 2008), with networks identified using a complete-linkage hierarchical clustering procedure (Sørensen, 1948) based on ROI-to-ROI anatomical proximity and functional similarity metrics (Nieto-Castanon, 2020). Results were thresholded using a combination of a p < 0.05 connection-level threshold and a familywise corrected p-FDR < 0.05 cluster-level threshold (Benjamini & Hochberg, 1995).

#### Graph theory

The graph measures of interest were estimated for the entire network of ROIs (N = 57), by using a fixed network cost level (i.e., keeping the strongest 15% of connections) to threshold the adjacency matrix. The graph theory measures were estimated for the positive adjacency matrix. For each GT measure of interest, a two-sided paired t-test was performed to examine the group-level changes from baseline at each time point. Results were thresholded using a familywise corrected p-FDR < 0.05 (Benjamini & Hochberg, 1995). To assess how individual ROIs change their contribution to the network structure and function, we used node degree and centrality to gain insights into both local connectivity and the role of specific ROIs within network dynamics.

Node centrality was defined as the global efficiency at a node which is computed as the average inverse-distances between this node and all other nodes in the graph. This measure of node centrality characterizes the degree of global connectedness of each ROI. Node centrality is useful for determining the relative importance of a node within the entire network. Centrality metrics help identify nodes that are not only well-connected but also crucial overall network dynamics. Node degree was defined at each node as the number of edges from/to each node. Node degree characterized the degree of local connectedness of each ROI within a graph. For example, a high degree indicates that a node is highly connected within the network, suggesting that it might play a significant role in local network interactions.

#### Overlap index

The overlap index was computed to assess how the overall direction of changes from baseline varies across subjects over time, without considering the magnitude of the effects. This approach allowed us to evaluate the temporal dynamics of individual differences to cocaine use, abstinence and re-exposure. For each subject, timepoint, and ROI, the change from baseline was coded as +1 for positive effects and-1 for negative effects. Subsequently, for each timepoint and area, the mean effect was computed across all subjects to obtain the overlap index. The overlap index ranges between 0, i.e., suggesting that there is a balance between positive and negative effects, and hence, no overall trend in one direction; and 1 indicating that all subjects exhibited positive/negative effects. Note that we do not differentiate between positive and negative effects, hence, the range of the index is 0 to 1.

### Behavior

We collected the following data: DA: 69 sessions; MO: 74 sessions; ME: 67 sessions; FA: 33 sessions (note: data collection was interrupted in this subject after 45 days of cocaine SA due to VAP complications). To quantify drug uptake and escalation of use, our outcome variable was daily amount of drug intake. To quantify drug seeking, our outcome variable was the average number of LPs during the drug unavailability period and the number of LPs during the drug availability period that were not rewarded with a cocaine infusion – i.e., LPs during the infusion time.

### Functional connectivity and behavior

To assess whether individual differences in behavior explain the variability in FC changes from baseline, we used the average node degree change from baseline for each condition and subject (i.e., Timepoint – Baseline). The behavioral measures were the unrewarded and rewarded LPs during the drug availability period and the TO LPs. The behavior was divided into 4 segments: SA before T1, SA between T1 and T2, SA after T2 until the end, and re-exposure. For each segment and NHP, the behavioral variables were summed across blocks and averaged across sessions. For each ROI, the resulting values were used as predictors in a multiple regression model with the change in node degree or centrality as dependent variables. We used permutation testing to assess the statistical significance of the observed relationships between behavioral predictors and changes in graph theoretical measures. To that end, we maintained the structure of the behavioral predictors constant while randomly permuting the dependent brain variable across the different runs. This process was repeated for 1000 iterations to generate a null distribution of the regression results, specifically the explained variance (R^2). By comparing the observed values to this permutation-derived null distribution, we calculated p-values to determine the likelihood that the observed relationships could occur purely by chance. Furthermore, we established a secondary threshold criterion across ROIs, whereby only ROIs exhibiting an R^2 value within the top 25th percentile of ROI models was deemed robust and indicative of significant predictive power.

## Supplementary material

**Supplementary Figure 1.**
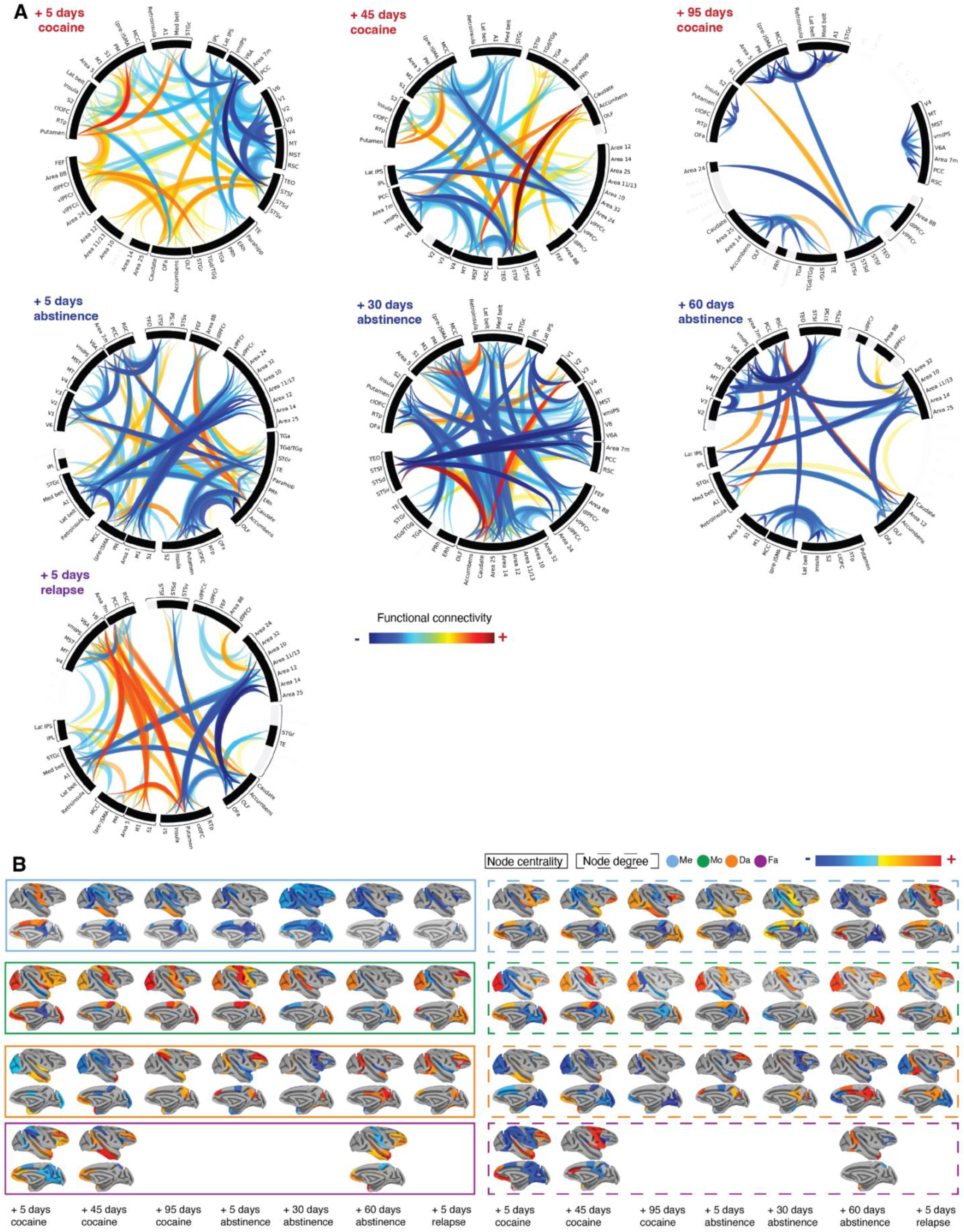
(A) Magnitude changes from baseline in functional connectivity. Multivariate parametric statistics with random-effects across subjects and sample covariance estimation across the time points were used to establish statistical significance. Inferences were made at the cluster level, which were defined as groups of similar connections - i.e., the clustering of the connections differs across time points according to their functional similarity and was integral to the statistical inferences. Results were thresholded using p < 0.05 connection-level threshold and a familywise corrected p-FDR < 0.05 cluster level threshold. **(B)** Network topology (i.e., node centrality and degree) changes from baseline at the single-subject level. At each subject and time point, network measures were computed for both the positive and negative adjacency matrices. For each graph theory measure and time point, a two-sided paired t-test was used to examine the group-level changes from baseline. Results were thresholded using p < 0.05.

---

**Supplementary Table 1.** Overview of networks and associated areas.

| Network | Areas |
| --- | --- |
| Reward Network (RN) | Area 25, Area 32, Area 10, Area 14, Areas 11/13, Area 12, Nucleus Accumbens |
| Habit Network (HN) | Caudate, Putamen |
| Salience Network (SAL) | Insula, Area 24, IPL |
| Default-mode or Self-directed Network (DMN) | PCC, RSC, Area 7 |
| Central Executive Network (CEN) | dIPFC, vIPFC |
| Dorsal Attention Network (DAN) | IPS, FEF |
| Somatomotor Network (SMN) | M1, PM, (pre-)SMA, S1, S2, MCC, Area 5, Retroinsula |
| Multimodal Integration Network (MIN) | STS, TGa, TGd/g |
| Medial Temporal Lobe Network (MTL) | Parahippocampal cortex, ERh, PRh |
| Auditory Network (AUD) | A1, Belt, STG |
| Visual Network (VIS) | MST, MT, V1, V2, V3, V6, V6A, TEO |

## Notes

**Acknowledgements** We thank Mark Thomas, Hank Jedema, Charles Bradberry as well as Melanie Graham for valuable discussions and for help with animal care and preparation. This work was supported by NIH grants R01 MH128177 (JZ), P30 DA048742 (AMG, SRH, AZ, JZ), and a UMN AIRP award (JZ, AZ).

### Competing Interest Statement

The authors have declared no competing interest.

